# Changes in cortical beta power predict motor control flexibility, not vigor

**DOI:** 10.1101/2025.01.23.634491

**Authors:** Emeline Pierrieau, Claire Dussard, Axel Plantey-Veux, Cloé Guerrini, Brian Lau, Léa Pillette, Nathalie George, Camille Jeunet-Kelway

## Abstract

The amplitude of beta-band activity (β power; 13-30 Hz) over motor cortical regions is used to assess and decode movement in clinical settings and brain-computer interfaces, as β power is often assumed to predict the strength of the brain’s motor output, or “vigor”. However, recent conflicting evidence challenges this assumption and underscores the need to clarify the relationship between β power and movement. In this study, sixty participants were trained to self-regulate β power using electroencephalography-based neurofeedback before performing different motor tasks. Results showed that β power modulations can impact different motor variables, or the same variables in opposite directions, depending on task constraints. Importantly, downregulation of β power was associated with better task performance regardless of whether performance implied increasing or decreasing motor vigor. These findings demonstrate that β power should be interpreted as a measure of motor flexibility, which underlies adaptation to environmental constraints, rather than vigor.

## Introduction

Movement is associated with a decrease in the power of the brain electrical signal in the beta frequency band (β; 13-30 Hz), which can be detected non-invasively with electroencephalography (EEG) over motor cortical regions^1^. This decrease in β power has long been thought to reflect the activation of motor cortical neurons^2^. It also occurs without actual movement during motor imagery, though in an attenuated way^3^. Thus, β power is widely used to decode movement-related activity during motor imagery in the field of brain-computer interfaces (BCIs)^4^. In addition, β power has been the target of numerous non-invasive and invasive neurostimulation studies aiming to restore motor function in several motor disorders, such as Parkinson’s disease^5,6,7^ and stroke^8,9^. However, these studies provided mixed evidence regarding the efficacy of modulating β power on improving motor function, probably due to a poor understanding of the precise influence of β power on movement^10,11,12,13^.

From an electrophysiological perspective, EEG spectral power is influenced by the degree of spatial and temporal synchrony in local activation of cortical neurons^14^. The extent of neuronal synchronization across different frequency bands is thought to underlie distinct functions. Previous evidence suggested that β power is associated with the strength of motor output or movement “vigor”^15,16,17^. This hypothesis is supported by non-invasive neurostimulation studies that showed significant decreases in movement speed^18^ and force^19^ when driving the activity of motor cortical neurons at a β rhythm. Moreover, recent studies showed that downregulation of β power with neurofeedback (NF), which consists of providing real-time feedback of brain activity,^20^ can speed up movement initiation^21,22,23^. According to this view, reducing β power leads to increased movement vigor. Yet, some studies failed to find significant correlations between changes in β power and reaction time (RT)^24,16^, or even to find any significant difference in β power between movements that were performed at very different speeds and forces^25,10,26,27^. These inconsistent findings led to an alternative hypothesis, according to which the impact of β power on motor function should be interpreted in terms of both efferent and afferent signal processing within motor cortical regions^28,29^. Indeed, decreasing β synchronization is thought to facilitate the processing of novel incoming inputs^30,31^. Hence, downregulation of β power could foster updating of the motor command and, thereby, improve adaptation to environmental constraints or motor “flexibility” ^32,33^. Such an assumption may reconcile the inconsistent findings mentioned above. If β power affects motor flexibility, one may expect a significant association between β power and RT only in tasks in which participants are asked to initiate their movements as fast as possible. Furthermore, according to this view, downregulation of β power might not only speed up movement initiation and execution, as formerly demonstrated^21,22,23,34^, but it may also slow down movements if this leads to better task performance. In other words, downregulation of β power may result in higher movement vigor (i.e., stronger/faster movements) only when it concurs with task constraints and allow better task performance. To our knowledge, such a hypothesis has not been properly tested yet.

The present study aimed to clarify the impact of motor cortical β power modulation on movement execution by assessing whether changes in β power rather predict motor vigor or flexibility. To do so, a total of 60 participants were trained to down- and up-regulate their motor cortical β power through visual NF. In a first experiment (EXP1), NF was followed by a force task, in order to determine if modulations of β power specifically impact the motor variable involved in task performance (i.e., force) or rather all motor variables related to movement vigor (i.e., force, RT and movement time (MT)). In a second experiment (EXP2), NF was followed by a speed task, in which participants were asked to move either at a fast or a slow pace. The goal of EXP2 was to determine if the direction of the relationship between β power and movement speed varies according to speed instruction (motor flexibility hypothesis) or if it consistently remains negative (motor vigor hypothesis).

## Results

A total of 60 participants were included in EXP1 (n = 30) and EXP2 (n = 30). Trials of EXP1 and EXP2 included a NF phase, followed by a motor task. In EXP1, participants executed an isometric motor task in which they were asked to squeeze a dynamometer with their right hand with a force superior or equal to 70% of their maximal force and to maintain it for 5 s. In EXP2, participants performed a dynamic motor task consisting of 4 repetitions of opening of the right hand, either at a fast (Fast), slow (Slow) or Comfortable (Comf) pace, determined on an individual basis. In EXP1 and EXP2, NF was designed in order to enforce downregulation of β power (β-down) in some blocks, and upregulation of β power (β-up) in other blocks. Blocks of passive control condition (Sham-Passive) were also included (see Figure 1 and Methods for details).

**Figure 1.**
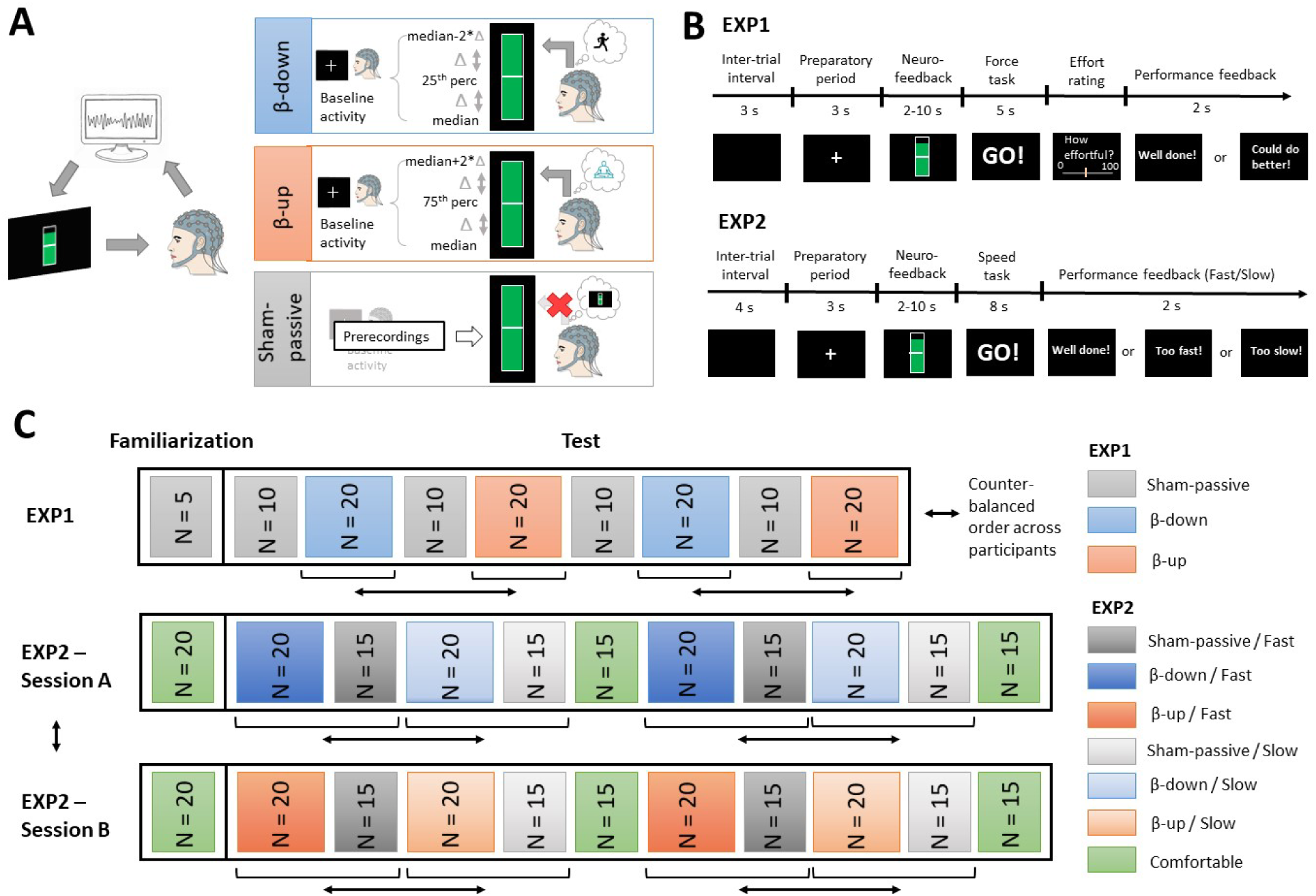
Experimental protocol of EXP1 and EXP2. **A.** Left, schematic illustration of NF. In EXP1 and EXP2, β power of the signal recorded over the left motor cortex with EEG was computed and represented as a virtual gauge. Participants were asked to try to fill up the gauge for as long as it was presented on the screen, which required them to self-regulate their β power. Right, illustration of the NF conditions that were implemented in EXP1 and EXP2. In β-down (blue) and β-up (orange), the level of the gauge increased as participants respectively decreased and increased their motor cortical β power in regard to baseline values that were predetermined individually. Participants were encouraged to use motor imagery and relaxation/selective attention to facilitate self-regulation of their motor cortical β power in β-down and β-up, respectively. In Sham-Passive (gray), the level of the gauge varied according to β power from prerecordings and participants were asked not to try controlling the level of the gauge. **B.** Top to bottom: name, duration and illustration of visual stimuli that were presented on screen during each phase of a trial in EXP1 (top) and EXP2 (bottom). **C.** Presentation order of blocks of trials in EXP1 (first row) and EXP2 (second and third rows). Blocks of trials are illustrated as colored rectangles in their order of presentation during the experiment, starting first with a block of familiarization trials. N refers to the number of trials included in each block. The color of the rectangles indicates the experimental condition that each block of trials pertains to (see legends on the right). The bidirectional black arrows indicate conditions that were counterbalanced across subjects (see Methods section for details).

### 1. NF condition (EXP1 and EXP2) and speed instruction (EXP2) significantly impacted motor cortical β power

The first step of data analysis consisted in determining if the NF paradigm used in EXP1 and EXP2 induced significant changes in motor cortical β power. Cluster-based permutation tests showed significant reduction of β power over a left central cluster of electrodes in β-down as compared to β-up, centered on C3 electrode in EXP1 (Figure 2A, top panel) and D19 (C3 equivalent in the ABCD system) electrode in EXP2 (Figure 2A, bottom panel). These electrodes were the ones used for NF computation (see Methods). Thus, NF condition (β-down versus -up) had a significant influence on motor cortical β power. There was not any significant cluster either for the comparison of β-down and Sham-Passive, nor for the comparison of β-up and Sham-Passive. Pooling data from both EXP1 and EXP2, an ANOVA on β power from C3/D19 electrode confirmed this result: NF conditions significantly impacted β power (F(1.6,88.6) = 4.7, p = 0.019), with a significant reduction of β power in β-down in comparison to β-up (W(57) = 510, p = 0.028, r = −0.40). The mean β power in Sham-Passive was intermediate between β-down and β-up and did not significantly differ from mean β power in β-down (t(57) = −1.6, p = 0.159, d = −0.22) and β-up (W(57) = 699, p = 0.227, r = 0.18) (Figure 2B). Within-subject comparisons of β power in β-down versus β-up indicated that on average 57% of participants (60.7% in EXP1, 53.3% in EXP2) downregulated their β power in β-down relative to β-up with a mean effect size (Cohen’s d) of −1.21 (SD = 0.51). Further analysis of online measurements of β power during NF trials confirmed that participants significantly decreased their β power in β-down as compared to β-up, and that they did so from the first block of each experiment (Figure S1). Inter-individual variability in modulations of β power between NF conditions was considered in the following analyses and used as a leverage for dissociating the respective effects of β power modulation and NF instruction on motor variables.

**Figure 2.**
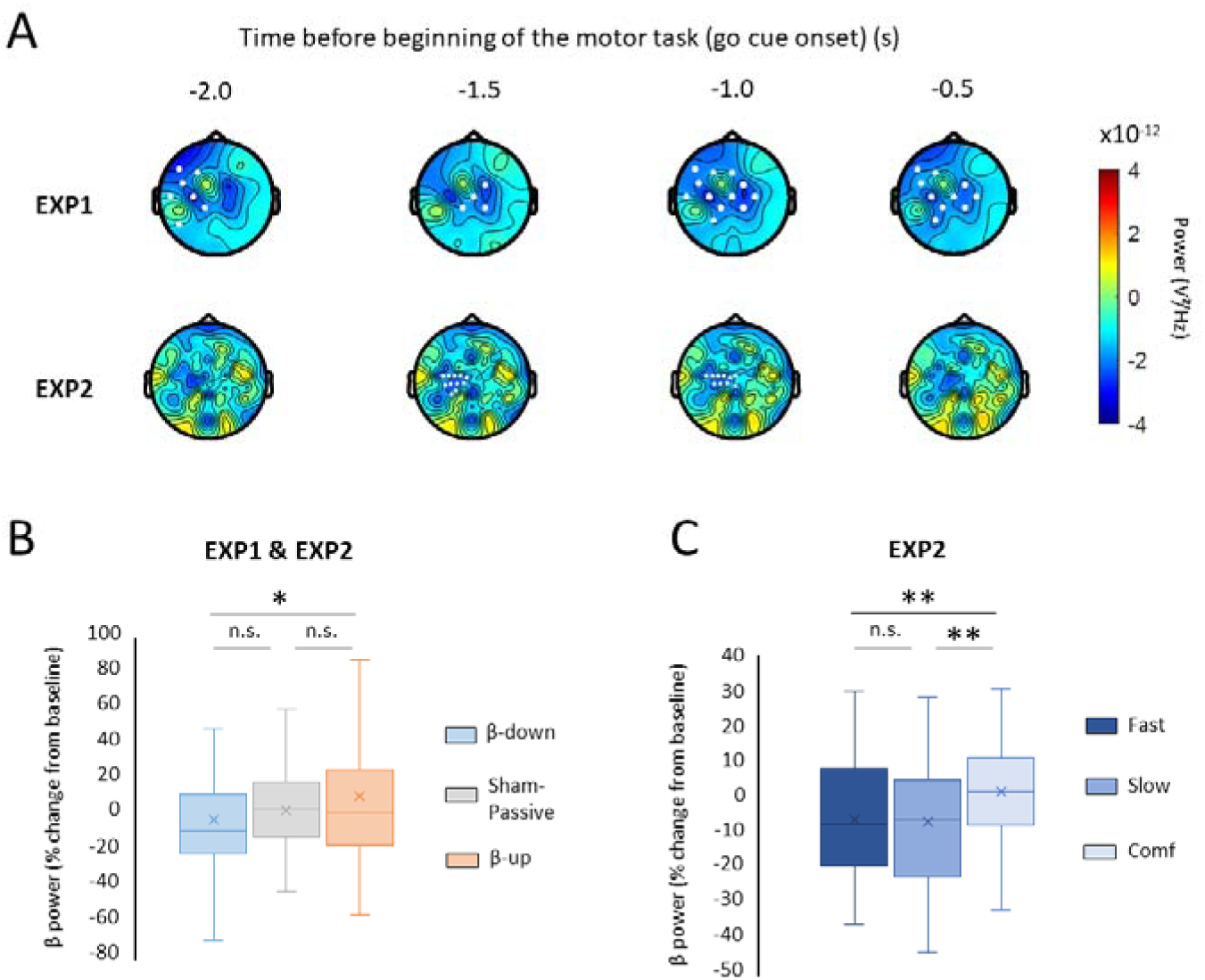
NF condition and speed instruction significantly impacted motor cortical β power. **A.** Topographical maps of mean β power difference between β-down and β-up at 4 time instants before go cue onset in EXP1 (top panel) and in EXP2 (bottom panel). The overall mean of the difference across subjects is represented for each experiment. Cold (blue) and warm (yellow to orange) colors illustrate respectively a decrease and an increase in β power in β-down as compared to β-up. Electrodes showing a significant difference in β power in β-down versus β-up are highlighted in white. **B, C.** Boxplots of average percentage of change in β power from baseline across NF conditions (EXP1 and EXP2 data pooled) and speed instruction (EXP2 data). Crosses indicate mean values. **p < 0.01, *p < 0.05, n.s. = not significant

The second step of the analyses consisted in assessing the effect of speed instruction on motor cortical β power in EXP2, since β power has been previously linked to movement speed^11,18,26,16^. Speed instruction (Fast, Slow, Comf) significantly influenced β power (F(1.6,47.2) = 6.6, p = 0.005). β power was higher in Comf in comparison to both Fast (t(29) = −2.5, p = 0.025, d = −0.46) and Slow (t(29) = - 3.3, p = 0.008, d = −0.60) conditions, but no significant difference in β power was found between Fast and Slow conditions (t(29) = 0.3, p = 0.734, d = 0.06) (Figure 2C). Speed instruction was also included as factor in the following analyses, which aimed at studying the relationship between β power changes and motor variables.

### 2. Modulations of motor cortical β power specifically impacted motor variables that determined task performance (EXP1)

Most NF studies targeting motor cortical β power reported a facilitatory effect of β power downregulation on RT and MT^21,22,23,34^, suggesting that reducing β power may increase movement vigor and, therefore, effort exertion. Yet, whether the reported boosting effect of downregulation of β power on RT and MT reflects an overall increase in movement vigor or rather an increase in motor flexibility, resulting in better task performance, remains unclear. In order to address this question, we designed a motor task where performance exclusively relied on movement force (EXP1, see Methods).

As a preliminary, we tested the correlation among motor variables. Between-subject correlation analyses confirmed that the mean force exerted during the motor task of EXP1 was neither significantly correlated with RT (r = 0.02, p = 0.938), nor with MT (r = 0.06, p = 0.758). RT was negatively correlated with MT (r = −0.59, p = 0.001). In this context, if changes in β power affect movement vigor, one should expect significant linear associations between β power on one hand and force, RT and MT on the other hand. Alternatively, if changes in β power affect motor flexibility, then one should expect a significant association between β power and force, but not between β power and RT or MT, as those variables were irrelevant to task performance and decorrelated from movement force (Figure 3A, 3D and 3G).

**Figure 3.**
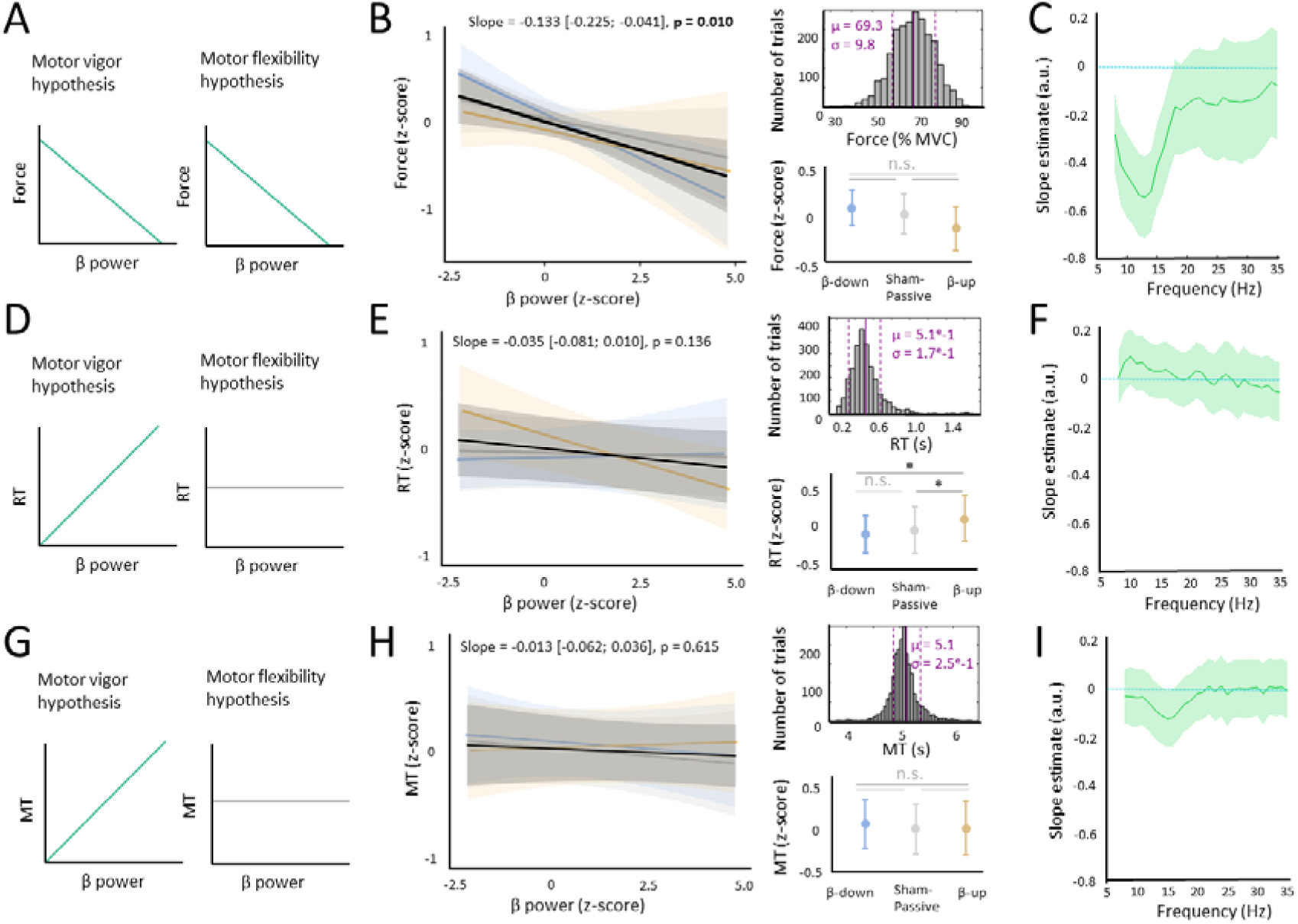
Modulations of motor cortical β power specifically impacted motor variables that determined task performance in EXP1. **A, D, G.** Schematic illustration of expected results according to the motor vigor (left) and motor flexibility (right) hypotheses. Green and gray curves indicate expected significant and non-significant associations, respectively. **B, E, H.** Left, linear regression lines of each motor variable tested (B: Force, E: RT, H: MT) on β power, estimated from LME models. Lines depicting results in β-down, β-up and Sham-Passive are respectively represented in blue, orange and gray. The main effect of β power (i.e., averaged across NF conditions) on motor variables is depicted as a black line. Shaded areas indicate 95% confidence intervals. Slope estimates of each line and their respective 95% confidence intervals are written on top of each plot, with p-value from one-sample t-test of mean slope against zero, written in bold when significant. Top right, distribution of single trial values of each motor variable. Mean (µ) and standard deviations (σ) of distributions are written on each panel in magenta for correspondence between z-scores used in the models and actual values. Bottom right, estimated marginal means of motor variables across NF conditions (β-down: blue, β-up: orange, Sham-Passive: gray). Error bars represent 95% confidence intervals. *p < 0.05, n.s. = not significant. **C, F, I.** Slope estimates of the linear regressions of motor variables (C: Force, F: RT, I: MT) on signal power at each 1 Hz frequency band between 8 and 35 Hz. In all plots, shaded areas illustrate 95% confidence intervals.

Linear mixed-effect (LME) models were conducted to determine if β power significantly predicted each motor variable (force, RT and MT) across trials and participants. NF instruction was included in these LME models to assess the respective effects of NF instruction and β power changes on motor variables. This approach enabled the dissociation of the influence of the different mental strategies that were applied across NF conditions (i.e., motor imagery, relaxation/selective attention and passive) from the impact of actual changes in β power during NF trials on motor variables. Participant ID was included as random factor and a full random effect structure was adopted to account for between-subject variability in the effects of interest (see Method for details). Separate LME models were conducted on mean force, RT and MT. The frequential selectivity of brain-behavior associations was further examined by running the same LME models but with power in 1 Hz bins from 8 to 35 Hz instead of β power.

Results from LME models showed a significant negative relationship between β power and movement force (F(1,20.6) = 8.1, p = 0.010) (Figure 3B, left). No significant effect of NF instruction was found on movement force (F(2,22.9) = 1.9, p = 0.178), nor any significant interaction between β power and NF instruction (F(2,12.1) = 1.4, p = 0.282) (Figure 3B, bottom right). Frequential selectivity analyses further demonstrated a significant negative linear association between spectral power and movement force between 8 and 17 Hz (mean slope estimate = −0.420, min = −0.540, max = −0.247) (Figure 3C). Converting z-scores back into actual values (Figure 3B, top right), this model predicted that each 1% increase in force was associated with a 8.1% decrease in low β power between 8 and 17 Hz.

In contrast, RT was significantly influenced by NF instruction (F(2,23.9) = 5.9, p = 0.008) (Figure 3E, bottom right), but not by β power (F(1,24.6) = 2.4, p = 0.136) (Figure 3E, left). RT was significantly increased in β-up in comparison to β-down (t(26.5) = 2.5, p = 0.030) and Sham-Passive (t(26.6) = 2.6, p = 0.049), whereas no significant difference in RT was found between β-down and Sham-Passive (t(26.5) = −0.5, p = 0.637). There was also a significant interaction between β power and NF instruction (F(2,43.9) = 3.2, p = 0.049). It reflected a negative linear association between β power and RT in β-up (slope parameter estimate = −0.106, 95% confidence interval = [-0.205; −0.007]), but this effect did not survive FDR correction (t(17.7) = −2.3, p = 0.110). No significant association was found between β power and RT in β-down and Sham-Passive (|t| < 0.3, p > 0.885). The lack of significant effect of β power on RT was not explained by the selected frequency band, as no significant association was found in frequential selectivity analyses between 8 and 35 Hz (Figure 3F).

Finally, there was not any significant effect of β power, NF instruction, or interaction between β power and NF instruction on MT (F < 0.7, p > 0.541) (Figure 3H). The analysis of frequential selectivity of the association between spectral power and MT showed a local negative linear relationship at 15 and 16 Hz (Figure 3I). The mean slope estimate for this effect (−0.125) was less pronounced than for the association found between low β (8-17 Hz) power and force (−0.420).

Overall, in EXP1, downregulation of β power was associated with increased force, without significantly affecting RT and MT. The relationship between β power and force was the strongest in the low β band (8-17 Hz). These findings support the view that motor cortical β power is associated with motor flexibility rather than with motor vigor. However, it may be argued that these two processes were not fully dissociated in EXP1 as motor flexibility relied on increased motor vigor in the form of higher force. Therefore, we designed a second task (EXP2) dissociating the impact of modulations of β power on motor flexibility and vigor by assessing the directionality of the relationship between β power and MT.

### 3. The direction of the relationship between β power and performance-related motor variables depended on task instruction (EXP2)

In EXP2, the motor task consisted in hand opening and closing at either fast or slow pace in separate trials (see Methods). In this context, if β power influences movement vigor, then downregulation of β power should speed up movements, reducing movement time (MT), regardless of the speed instruction of the motor task. In contrast, if β power impacts motor flexibility, then downregulation of β power should lead to shorter MT when instructed to move fast and, conversely, longer MT when instructed to move slowly. Hence, EXP2 paradigm aimed to further disentangle the motor vigor and flexibility hypothesis by confronting their opposite predictions in the slow movement condition (i.e., negative vs positive linear relationship between β power and MT according to motor vigor and motor flexibility hypotheses, respectively) (Figure 4A, 4D, 4G).

**Figure 4.**
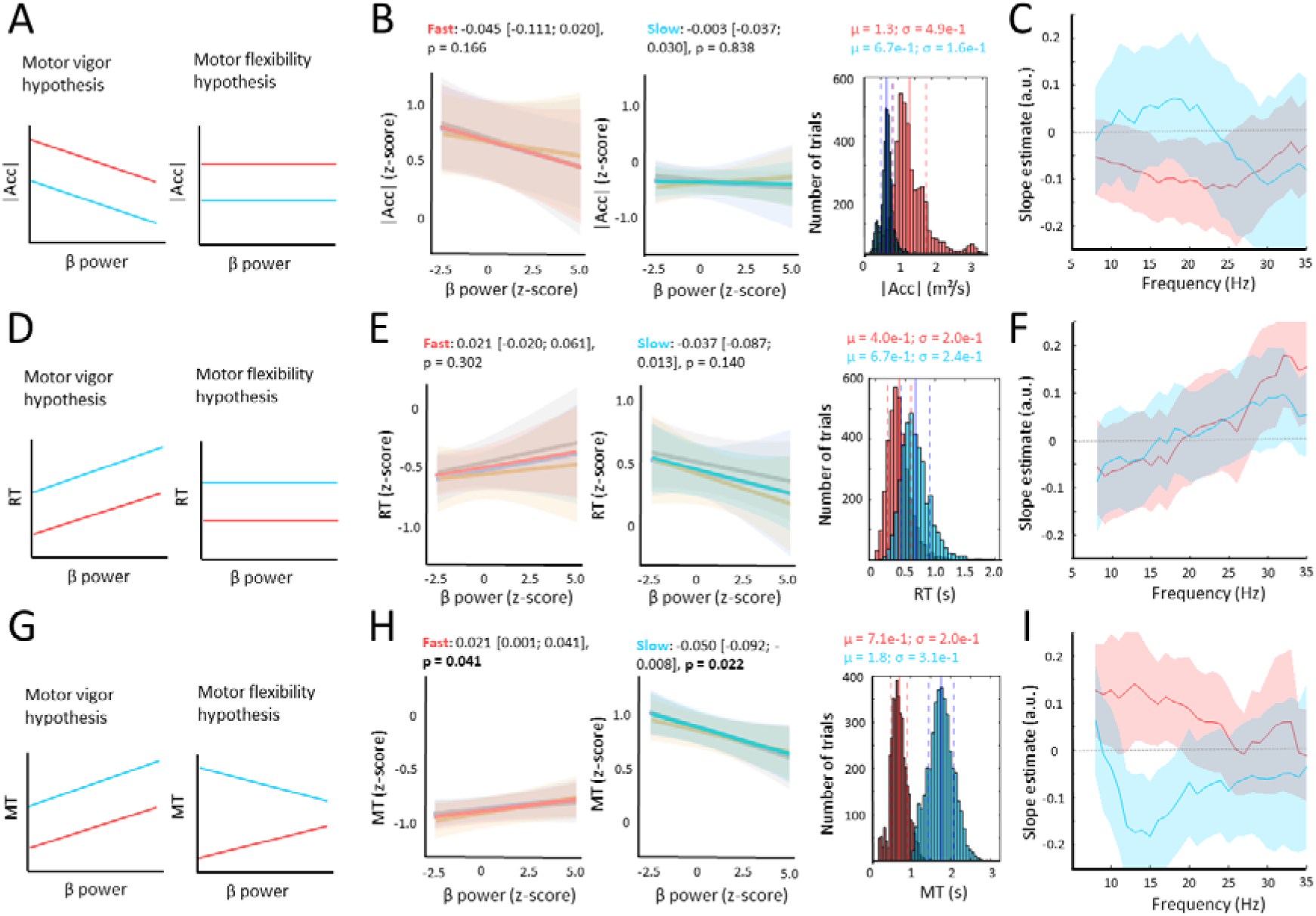
The direction of the relationship between β power and performance-related motor variables was reversed depending on task instruction in EXP2. **A, D, G.** Schematic illustration of expected results according to the motor vigor (left) and motor flexibility (right) hypotheses. Red and cyan curves indicate expected linear associations in Fast and Slow, respectively. **B, E, H.** Left, linear regression lines from LME models between β power and each motor variable tested (B: |Acc|, E: RT, H: MT) and NF condition (blue: β-down, orange: β-up, gray: Sham-Passive) and averaged across NF conditions in Fast (left plot, red line) and Slow (right plot, cyan line). Slope values of each line and their respective 95% confidence intervals are written on top of each plot, with p-value from one-sample t-test of mean slope against zero. Right, distribution of single trial values of each motor variable. Fast trials are represented in red and Slow trials in cyan. Mean (µ) and standard deviations (σ) of distributions are written on the right of each panel in Fast (red) and Slow (cyan) separately. **C, F, I.** Slope estimates of the association found between signal power and motor variables in LME models at each 1 Hz frequency band between 8 and 35 Hz. Red and cyan curves represent results in Fast and Slow, respectively. In all plots, shaded areas illustrate 95% confidence intervals.

The motor variables in EXP2 were MT, RT and mean absolute acceleration (|Acc|). Indeed, as EXP2 was a dynamic motor task, |Acc| was computed as a measure of vigor, in lieu of force (used in the isometric task of EXP1), considering that |Acc| has been used previously to measure motor effort in speed tasks^35^. The analysis of the correlation among motor variables showed that MT was significantly positively correlated with RT in Slow (r = 0.42, p = 0.021), but not in Fast movement conditions (r = 0.32, p = 0.085). MT and |Acc| were strongly negatively correlated in Fast (r = −0.90, p = 10^-11^) and, to a smaller extent, in Slow (r = −0.38, p = 0.040). Correlations between RT and |Acc| in Fast and Slow were not statistically significant (Fast: r = −0.36, p = 0.052; Slow: r = −0.29, p = 0.115). Notably, these correlations were expected in the context of this speed task and did not undermine testing EXP2 hypotheses. Indeed, unlike EXP1, the objective of EXP2 was not to determine if β power specifically predicts one motor variable, but rather the impact of task instruction (Fast vs Slow) on the relationship between β power and a performance-related variable (MT).

As for EXP1, we used linear mixed-effect (LME) models to test the respective effects of β power changes and NF instruction on each motor variable. Speed instruction (Fast, Slow) was also included as fixed-effect factor in the models. Separate LME models were conducted on |Acc|, RT and MT. The frequential selectivity of brain-behavior associations was further examined by running the same LME models but with power in 1 Hz bins from 8 to 35 Hz.

LME models confirmed that |Acc| was significantly impacted by speed instruction (F(1,29.0) = 53.4, p = 10^-8^). There was not any significant main effect of either β power (F(1,24.1) = 3.5, p = 0.073) or NF instruction (F(2,27.2) = 1.0, p = 0.380), and the interaction between speed instruction and β power was not significant (F(1,26.3) = 0.9, p = 0.340) (Figure 4B). The frequential selectivity analysis showed a negative linear relationship between |Acc| and β power between 16 and 24 Hz in Fast (mean slope = −0.108, min = −0.098, max = −0.120) (Figure 4C). No significant association was found between |Acc| and the power of any of the tested frequency bands in Slow.

Likewise, RT was significantly impacted by speed instruction (F(1,28.1) = 221.2, p = 10^-14^), but not by β power and NF instruction (F < 1.5, p > 0.261) (Figure 4E). There was a significant interaction between β power and speed instruction (F(1,20.3) = 4.8, p = 0.041). However, the analysis of simple effects of β power according to speed instruction did not show any significant linear association between RT and β power either in Fast or in Slow (Figure 4E). In Fast, a positive linear relationship between RT and power between 29 and 35 Hz (mean slope = 0.150, min = 0.122, max = 0.179) (Figure 4D). No significant association was found between RT and signal power in any of the frequency bands tested in Slow.

Results from the LME model confirmed that MT was significantly influenced by speed instruction (F(1,28.9) = 563.6, p < 10^-16^). There was not any significant main effect of either β power or NF instruction (F < 2.1, p > 0.165). Importantly, a significant interaction was found between β power and speed instruction (F(1,27.9) = 8.1, p = 0.008). Post-hoc analysis showed that the direction of the relationship between β power and MT was reversed depending on the speed instruction: it was positive in Fast (t(23.1) = 2.2, p = 0.041) and negative in Slow (t(28.3) = −2.4, p = 0.022) (Figure 4H). Frequential selectivity analyses showed that, in Fast, signal power was positively associated with MT in the low alpha band (8-10 Hz), as well as in the low β band (13-18 Hz) (mean slope = 0.119, min = 0.098, max = 0.141) (Figure 4I). Based on this model, each 15.38% decrease in low alpha/low β power predicted a 10 ms decrease in MT in Fast. In Slow, there was a negative linear relationship between MT and signal power between 12 and 18 Hz (mean slope = −0.156, min = −0.121, max = −0.181) (Figure 4I). Based on this model, each 7.13% decrease in low β power predicted a 10 ms increase in MT in Slow.

In sum, in EXP2, modulations of β power impacted MT in a task instruction-dependent manner: reducing β power resulted in faster movements when participants were instructed to move fast and, conversely, it resulted in slower movements when participants were instructed to move slowly. The association between motor cortical power and MT was the strongest in the low β band, both in Fast and Slow conditions.

## Discussion

The aim of the present study was to dissociate the impact of changes in motor cortical β power on movement vigor and motor flexibility, in order to clarify how motor cortical β power relates to movement. This was achieved through two experiments (EXP1 and EXP2) combining a NF task to train self-regulation of motor cortical β power with a motor task, in order to assess the effects of β power down- and up-regulation on movement execution. EXP1 findings showed that modulations of β power were significantly associated with changes in force, but not with changes in RT and MT, in the context of a force task. More precisely, reducing β power resulted in higher force exertion. Therefore, changes in β power appeared to specifically impact the motor variable that was involved in task performance. This result was replicated and extended in EXP2, in which modulations of β power were significantly associated with MT task instruction-dependent manner in the context of a speed task. EXP2 results highlighted that the direction of the relationship between β power and MT depended on task instruction: β power downregulation was associated with faster movement (i.e., decreased MT) when instructed to move at a fast pace vs slower movement (i.e., increased MT) when instructed to move at a slow pace. Finally, no significant influence of NF instruction was found on any of the motor variables tested except for RT in EXP1. In both experiments, the association between β power and performance-related variables was the strongest in the low β band.

Overall, the present findings show that changes in β power do not appear to specifically impact one motor variable, but rather any motor variable that is involved in task performance. While most NF studies reported a shortening of RT accompanying NF-induced decreases in β power^21,22,23^, results of EXP1 and EXP2, gathered on a total of 60 participants, did not show any significant effect of modulations of β power on RT. Nevertheless, a significant linear association was found between β power and force in the force task of EXP1, as well as between β power and MT in the speed task of EXP2. These associations appeared task-specific as there was not any significant linear association between β power and MT in EXP1, nor between β power and |Acc| -which can be used as a measure of motor effort in speed tasks^35^- in EXP2. It is only when breaking down β power into 1 Hz bins ranging from 8 to 35 Hz that a significant negative association between 15-16 Hz power and MT in EXP1 was found, as well as between 16-24 Hz power and |Acc| in Fast in EXP2. Still, the former was considerably weaker than the association found between low β power and force (slope estimate was almost 4 times lower), and the latter was likely explained by the strong correlation (r = −0.90) found between MT and |Acc| in Fast.

At first glance, the significant negative relationship found between is β power and force in EXP1 could be interpreted as downregulation of motor cortical β power boosting activation of motor regions, which has been proposed to increase movement “vigor” and effort exertion, resulting in faster and stronger movements^15,16,17^. However, both EXP1 and EXP2 findings invalidate such a hypothesis. Indeed, in EXP1, RT and MT were unaffected by changes in β power, showing that downregulation of β power was associated with stronger, but not faster movements. Even more strikingly, EXP2 results show a reversal of the direction of the relationship between β power and MT with speed instruction, with downregulation of β power resulting in faster movements only when participants were instructed to move fast. Conversely, decreasing β power was associated with longer MT when instructed to move slowly such that, in either way, downregulation of β power enhanced task performance. According to this result, the effect of changes in β power on motor behavior that has been consistently reported by previous studies (i.e., faster movement initiation and execution when reducing β power) is unlikely to be explained by changes in movement vigor. Rather, it supports the view that β power is associated with motor control flexibility. Indeed, decreasing neuronal synchronization in the β band (i.e., decreasing β power) is thought to facilitate the processing of novel incoming inputs ^13,30,31^. In the context of motor planning, it implies that motor commands are more readily actuated based on environmental disturbances, resulting in more flexible motor control for better adaptation. In line with this hypothesis, several studies demonstrated significant reduction of β power during early phase of visuomotor adaptation tasks, which attenuates once errors decrease and participants no longer need to adapt their behavior^36,37,33^. In EXP2, both fast and slow movement conditions required a certain degree of motor adaptation as participants needed to adjust their movement speed to an instructed MT, which was 3 standard deviations away from their habitual, comfortable MT. Results confirmed that β power was significantly lower for fast and slow movements in comparison to movements executed at a comfortable pace. The significance of these findings may extend to pathological brain states, as they are consistent with recent evidence on bradykinesia in Parkinson’s disease, which is associated with excessive β synchrony within motor cortex-basal ganglia loops^11^. Bradykinesia has been suggested to originate from a deficit in switching between dynamic and stable movement states, leading to less flexible or “rigid” motor control, rather than from a deficit in vigor^38^. In line with the motor flexibility hypothesis, a recent study demonstrated that reduction of β power at the subcortical level with electrical stimulation of the subthalamic nucleus can lead to larger absolute adjustments of force irrespective of direction, and that the more the stimulated area was connected to the cortex, the greater the force adjustment^39^. Overall, the present study provides novel direct evidence of motor cortical β power as a relevant target for enhancing motor flexibility and, thereby, task performance.

Whether NF constitutes an adequate tool to reliably and sustainably enhance motor flexibility through downregulation of β power remains an open question. Around 60% (34/58) of participants showed significant reduction of their β power in β-down in comparison to β-up, which is consistent with the proportion of responders typically reported in NF and BCI studies^40^. Additionally, the proportion of responders was possibly reduced in the present study due to the alternation between opposite trainings of self-regulation of β power (down- vs up-regulation) in the two experiments, which may have interfered with each learning process. Indeed, achieving self-regulation of specific patterns of brain activity with NF training relies on brain plasticity mechanisms that often require several days of training^41^. Providing longer uninterrupted NF training over several sessions and assessing motor performance pre- and post-NF training in each session represents a relevant alternative. In any case, optimizing β power regulation appears as a key factor to significantly impact motor flexibility with NF, considering that β power, and not NF instruction, accounted for motor effects in LME models. Still, NF instruction as such (that is, the mental strategies used during NF trials) may also induce non-specific effect, as demonstrated by the significant association between RT and NF instruction in EXP1. Yet, this effect was not replicated in EXP2 and, therefore, was distinct from the effect of β power down-versus up-regulation. Moreover, in the present study, a bidirectional NF was used in order to produce experimental conditions with significantly different level of β power before initiating movements (β-down and β-up) and overall maximize intra-individual variations in pre-movement β power. This manipulation enabled optimal feeding of predictive models of motor variables based on β power. Considering that NF was specifically used as a tool to determine the impact of modulations of β power on movement execution, the potential efficacy of NF training for improving motor performance should not be inferred from the present results. A proper sham condition (with active motor imagery) would be required to do so. Rather, as mentioned above, the present findings emphasize the importance of optimizing NF training to induce a reliable and strong enough decrease in β power from its baseline level, so that it may result in significant improvement of motor flexibility and, hence, task performance.

Finally, the task-specific associations found between β power and motor variables shed light on the importance of considering task constraints for β power-based applications such as BCIs. Indeed, β power is commonly used as input for machine learning algorithms to decode movement intention in motor imagery BCIs^4^. However, real time decoding of movement intention based on this method, especially using non-invasive signals such as EEG, remains challenging^42^. Here we show that environmental constraints critically matter when trying to decode movement intention based on motor cortical β power. What motor variables and how they can be decoded from changes in β power appear to strongly relies on the environmental constraints the subject is being exposed to. Hence, decoding of β power is likely relative rather than absolute. This is a critical point considering that, based on the models from the present study, similar β power values can be associated with substantially different movement kinematics. For instance, discriminating fast from slow movements may be complicated by similar β power values reached in the two conditions. While the development of more elaborate and precise machine learning algorithms might help overcoming this issue^43^, feeding those algorithms with other EEG data than motor cortical β power may be key to improve BCI performance. A more thorough understanding of the dynamics of EEG signal underlying movement preparation and execution appears crucial to this aim.

## Methods

### Participants

Thirty individuals (15 females, age (mean ± standard deviation): 22 ± 3 years old) participated in EXP1 and thirty other individuals (15 females, age (mean ± standard deviation): 24 ± 4 years old) participated in EXP2. Inclusion criteria included being right-handed or ambidextrous (the overall criterion being that they would perform daily motor tasks with their right hand), free of any known neurological or psychiatric condition and of any recent injury to the right upper limb that could limit their movements or cause any pain, and having normal or corrected-to-normal vision. In EXP1, data from two participants were removed from the analyses because of excessive noise in their EEG data. All participants gave their written informed consent before participation in the study, which had been approved by the French committee for the protection of individuals (CPP number 2022-A00626-37). This study conformed to the standards set by the latest version of the Declaration of Helsinki.

### Material

#### 1. EXP1

Participants were first seated in front of a computer screen (60 x 34 cm) that displayed the visual stimuli used for the experiments. A 32-channel EEG system was used (EEGo Sports, ANTneuro) with a 500 Hz acquisition rate. Participants were asked to hold a dynamometer (K-Force Grip, Kinvent) with their right hand without exerting any pressure on it unless receiving the instruction to do so. Data from the dynamometer was acquired using Bluetooth® and custom scripts in Unity 2021.3.15f1, with a 75 Hz acquisition rate. Acquisition of EEG data was conducted with OpenViBE 3.4.0^44^. Visual stimuli (including NF) were designed and displayed on screen using Unity 2021.3.15f1. Recordings from the dynamometer and EEG were synchronized by means of digital stimulation codes indicating the onset of each visual stimulus that were generated by OpenViBE and sent to Unity using Lab Streaming Layer (LSL).

#### 2. EXP2

Participants were first seated in front of a screen (60 x 34 cm) that displayed the visual stimuli used for the experiments. A 128-channel EEG system was used (ActiveTwo, Biosemi) with a 2048 Hz acquisition rate. A light sensor (LUX, BITalino) and an accelerometer (ACC, BITalino) were fixed on the distal phalange of the index of participants’ right hand, respectively on the ventral and dorsal side. A light was placed 20 cm above the hand. Participants were asked to keep the back of their right hand against the table, their palm facing the light. With this setting, the light sensor captured most light when the hand was fully open, and the least when the hand was closed. Data from the light sensor were acquired using Bluetooth®, OpenSignals 2.2.1 (PLUX) and custom scripts in Matlab R2023a (MathWorks), with a 1000 Hz acquisition rate. Acquisition of EEG data was conducted with OpenViBE 3.4.0^44^. Visual stimuli (including NF) were designed and displayed on screen using Psychtoolbox-3^45^ in Matlab. Recordings with EEG, the light sensor and the accelerometer were all synchronized by means of digital stimulation codes indicating the onset of each visual stimulus that were generated by OpenViBE and sent to Matlab using LSL.

### Overview of the experimental tasks

EXP1 and EXP2 included a bidirectional NF paradigm. Bidirectional NF consists of training participants to self-regulate a specific brain activity pattern (herein, motor cortical β power) in opposite directions (i.e., down- and up-regulation) in separate experimental blocks (β-down and β-up). β-down and β-up trials were performed within a single session in EXP1, and in 2 separate sessions in EXP2 (Figure 1C). The aim of this bidirectional NF paradigm was to maximize differences in β power between NF conditions (β-down and β-up), in order to properly assess the influence of β power on motor variables. Bidirectional NF indeed represents a more appropriate control than sham to determine brain-behavior relationships^46^. An additional Sham-Passive (control) condition was implemented, which included similar sensory inputs as in β-down and β-up, but without congruent NF nor mental strategy applied. The interest of this control condition was to measure β power and motor variables without active modulation of β power. This led to 3 NF conditions: active NF aiming to down-regulate motor cortical β power (β-down), active NF aiming to up-regulate motor cortical β power (β-up), and passive and sham NF aiming not to significantly modulate motor cortical β power (Sham-Passive) (Figure 1A).

#### 1. Trial timeline

##### a. EXP1

Trial timeline is depicted in Figure 1B. Each trial started with a black screen which lasted 3 s. Then, a white fixation cross appeared for 3 s, informing participants that they should stay still and look at the cross in preparation for the upcoming NF phase. The cross was then replaced by the NF, which consisted in a virtual gauge whose level varied according to online changes in motor cortical β power that were recorded with the EEG (see Neurofeedback section). The NF phase lasted between 2 to 10 s. It was immediately followed by the appearance of a go cue (“GO!”) indicating the beginning of the motor task, during which participants were asked to squeeze a dynamometer for 5 s at an individualized force level (see Motor task section). Once the motor task was over, participants rated their subjective perception of effort about the motor task on a virtual scale ranging from 0 to 100, by displacing a cursor with a computer mouse. Then, they received a feedback about their performance at the motor task for 2 s. Participants were asked to minimize eye and head movements during the presentation of the virtual stimuli, except during the intertrial interval (3-s black screen).

##### b. EXP2

Each trial started with a black screen which lasted 4 s. Then, a white fixation cross appeared for 3 s, informing participants that they should stay still and look at the cross in preparation for the upcoming NF phase. The cross was then replaced by the NF phase, which was identical to EXP1. It was immediately followed by the appearance of a go cue indicating the beginning of the motor task and speed instruction (drawing of a hand with either “FAST”, “SLOW” or “COMFORTABLE”), during which participants were asked to execute 4 repetitions of hand opening/closing, starting from an open hand position (see Motor task section). Once the motor task was over, participants received a feedback about their performance at the motor task for 2 s. Participants were asked to minimize eye and head movements during the presentation of the virtual stimuli, except during the intertrial interval (4-s black screen). Trials in which movements were performed at a comfortable pace did not include a NF phase nor feedback about performance at the motor task, as the aim of those trials was to obtain a baseline measurement of MT, without instruction nor active modulation of brain activity.

#### 2. Experiment timeline

##### a. EXP1

The session began with NF and force calibration. NF calibration consisted of a 1 min resting-state period to determine an individualized baseline β power value that was then used for NF settings (see Neurofeedback section). During this period, a white fixation cross was presented at the center of the screen. Participants were asked to keep their eyes open and looking at the fixation cross, while minimizing head, eye and body movements. They were also asked to place their right hand on the dynamometer without exerting any force on it. The cross was displayed for 1 min but only the last 30 s were used to compute baseline β power in order to ensure that β power was measured while participants were resting. Participants were then asked to squeeze the dynamometer with their right hand as hard as possible during 3 s in order to establish their maximal voluntary contraction (MVC) which value would be used to calibrate the motor task (see Motor task section). This procedure was repeated 3 times. MVC was calculated as the mean force of the 3 repetitions.

Afterwards, a first block of 5 Sham-Passive trials was presented to familiarize participants with the task. These familiarization trials were not included in the analyses. Then, participants performed 2 blocks of 20 β-down trials, 2 blocks of 20 β-up trials, and 4 blocks of 10 Sham-Passive trials, presented in an alternated order (Figure 1C, top). The interest in alternating presentation of NF conditions was to account for the effect of cognitive and physical fatigue, which ineluctably increased over the course of the experimental session. The presentation order of β-down and β-up conditions was counterbalanced across participants (either β-down/ β-up/ β-down/ β-up or β-up/ β-down/ β-up/ β-down). Each block of β-down or β-up trials was preceded by a block of Sham-Passive trials.

##### b. EXP2

In EXP2, blocks of β-down and β-up trials were performed in two separate sessions, taking place 1 to 3 weeks apart from each other. The organization of the blocks was similar between the two sessions (Figure 1C, bottom). The order of presentation of NF conditions (session 1 = β-down and session 2 = β-up or vice-versa) was counterbalanced across participants. Participants began each session with a calibration recording, consisting of one block of 20 trials without NF in which they were asked to execute the motor task (i.e., 4 repetitions of hand opening) at a comfortable pace. MT criteria for the motor task were set based on mean MT during this first block (see Motor task section). NF settings were based on β power during the preparatory period of this first block (see Neurofeedback section). Similar blocks (i.e., without NF and with movements executed at a comfortable pace) including 15 trials were presented at the middle (Block 7) and at the end of the experiment (Block 12). After the first block, a block of 10 familiarization trials with active NF (β-down or β-up depending on the session) was presented. Participants were not only familiarized with NF but also with the motor task, as this block included 5 trials with the motor task executed at a fast pace (Fast) and the 5 other trials at a slow pace (Slow). As in EXP1, familiarization trials were not included in the analyses. Afterwards, participants performed 4 blocks of 20 trials of β-down or β-up trials and 4 blocks of 15 Sham-passive trials, in an alternated order. Blocks of β-down or β-up trials were followed by blocks of Sham-passive trials with the same speed instruction (e.g., a block of β-down/Fast trials was followed by a block of Sham-Passive/Fast trials). Two different sequences of speed instructions could be presented across blocks: either Fast-Fast-Slow-Slow-Comf-Slow-Slow-Fast-Fast-Comf or Slow-Slow-Fast-Fast-Comf-Fast-Fast-Slow-Slow-Comf. These sequences were chosen such that one of the 2 blocks aiming to measure participants’ baseline MT (no NF, movements executed at a comfortable pace) occurred after a Fast block, and the other after a Slow block, thus accounting for potential changes in speed due to the influence of the speed instruction of the previous block. The order of presentation of speed instructions was counterbalanced across participants, but not across sessions (i.e., if one participant began with a Fast block in the first session, they would start with a Fast block in the second session as well).

### Neurofeedback

NF was represented as a virtual gauge. The level of the gauge reflected online changes of β power with respect to baseline β power. In EXP1, baseline β power corresponded to the median of the distribution of β power values during the 30-s resting-state recording. More precisely, online β power was computed as the squared amplitude of the signal recorded at C3 electrode, with bandpass (15-25 Hz, 4^th^ order Butterworth) and Laplacian (spatial coefficients: 6 for C3, −1 for adjacent electrodes (F3, FC5, CP5, P3, CP1, FC1)) filtering applied. Laplacian filtering was used to minimize contamination of the signal by artefactual sources such as electromyography by reducing the relative contribution of distant sources^47,48^. Bandpass filtering was applied on a narrower frequency band than the general β band (15-25 instead of 13-30 Hz) considering filter roll-off, that is the incomplete attenuation of frequencies beyond cutoff points^49^. Additionally, most studies reporting a significant impact of modulating β power on motor behavior targeted frequencies at or around 20 Hz^18,19,23^. Online β power was averaged over 500-ms epochs, with a 250-ms step, based on the method of He and colleagues (2020)^23^. The 250 ms overlap between consecutive epochs was introduced in order to smooth the signal and thus improve readability of the NF. Median, 25^th^ and 75^th^ percentiles of β power during the 30-s resting-state recording were determined based on a distribution of 120 online β power values (1 value per 250 ms) after outliers rejection. Outliers were identified as values inferior to median β power – 3*median absolute deviation (MAD) and superior to median β power + 3*MAD^50^. Median, 25^th^ and 75^th^ percentiles of baseline β power were then used for setting the NF individually (see paragraph below). In EXP2, baseline β power was computed during the 3-s preparatory period (white fixation cross, no NF and no movement) of the first block, representing 60 s of recording (3 s x 20 trials). The same method as in EXP1 was applied to calculate online β power, except that the Laplacian filter was applied on different but equivalent electrodes than the ones used in EXP1 (spatial coefficients: 6 for D19 (C3 equivalent), −1 for C24, D4, D10, D16, D26, A6). As in EXP1, median, 25^th^ and 75^th^ percentiles of β power during the baseline recording were computed and used for NF settings.

The design of the NF was the same in EXP1 and EXP2. A gauge, consisting of a white vertical rectangle cut by a white horizontal midline, was presented. The rectangle was filled with a bar, which height was determined by the relative difference between online and baseline β power. In β-down and β-up, the inferior boundary of the gauge corresponded to the median value of baseline β power. In β-down, the bar reached the horizontal midline (i.e., middle of the gauge) when online β power was equal to the 25^th^ percentile of baseline β power, whereas in β-up, the bar reached the horizontal midline when online β power was equal to the 75^th^ percentile of baseline β power. The absolute value of the difference between the median and the value at the horizontal midline (i.e., 25^th^ percentile in β-down, 75^th^ percentile in β-up) was computed as Δ (Figure 1A). The top of the gauge was reached when online β power was equal or inferior to median – 2*Δ in β-down, and when online β power was equal or superior to median + 2*Δ in β-up. Online β power was converted into the level of the gauge (i.e., height of the vertical bar) by normalizing its value between 0 and 1, 0 corresponding to the inferior boundary of the gauge (i.e., median baseline β power) and 1 to the superior boundary of the gauge (i.e., – 2*Δ in β-down, + 2*Δ in β-up). The vertical bar was filled in red when 0 < normalized online β power value < 0.5 (i.e., the level of the gauge was between the inferior boundary and the horizontal midline) whereas it was filled in green when normalized online β power value ≥ 0.5 (i.e., the level of the gauge was at the level of or above horizontal midline). Participants were asked to try to keep the gauge green as long as possible while it was displayed on the screen. Thus, participants were trained to maintain their online β power equal or inferior to the 25^th^ percentile of their baseline β power in β-down, and equal or superior to the 75^th^ percentile of their baseline β power in β-up. Different values (25^th^ and 75^th^ percentiles) were targeted in β-down and β-up in order to maximize the difference in online β power that participants could reach between the two conditions. Although different, these values were both extracted from the distribution of baseline β power values to make sure that they were attainable.

NF stopped and was replaced by the go cue as soon as participants maintained the gauge green for 2 consecutive seconds, or after 10 s if they failed doing so. This variable NF duration was chosen to increase the likelihood that online β power would be comprised in the targeted values at the time of go cue onset, in addition to further incentivize modulation of β power by giving participants the opportunity to reduce the duration of the experiment by improving their NF performance. In Sham-Passive, the design and settings of the gauge were the same as in β-down and β-up, the difference being that the level of the gauge was based on EEG prerecordings. In EXP1, prerecordings were extracted from pilot data and presented in the same order for all participants. In EXP2, prerecordings were replays of the participants’ online β power in the preceding block. Participants were asked if they noticed anything particular with the gauge in Sham-Passive trials once the second session was over, but none of them reported having noticed that it was based on replays of the preceding blocks.

Mental strategies that could be used to facilitate self-regulation of their motor cortical β power were explained to the participants before beginning the experiment. In β-down, participants were advised to use motor imagery, that is imagining themselves making a movement without actually executing it. Indeed, motor imagery can significantly reduce motor cortical β power^51,52^. When combined with NF, the reduction in β power induced by motor imagery can reach and even exceed the reduction in β power observed during motor execution^53^. Thus, motor imagery is commonly used as a mental strategy in NF studies intending to attenuate motor β power^23,34,54^. In opposition to motor imagery, relaxing body parts, conscious breathing and task-unrelated thinking appear as the most efficient strategies for increasing β power^55^. These strategies were detailed to the participants to help them improving their NF performance in β-up. Nonetheless, participants were told that making use of these mental strategies was not mandatory, and that they were free to use any other strategy that they would find efficient to improve their NF performance. The only restriction was to not make any body, head or eye (including closing/opening eyes) movements. Participants were also asked not to use any mental strategy in Sham-passive trials. They were not informed that the NF was a sham, but they were asked to merely keep their gaze on the gauge without trying to control its level, and to let their thoughts flow freely while staying focused during the motor task. The suggested mental strategies associated with each NF condition were reminded to the participants at the beginning of each block, both orally by the experimenter and with a message that was displayed for 5 s on the screen.

### Motor task

#### 1. EXP1

Participants were instructed to hold a dynamometer in their right hand and to start squeezing it as soon as the go cue appeared on the screen. They were asked to try to maintain the force they exerted above 70% of their MVC, as long as the go cue stayed on (5 s). This force threshold was selected to keep the task challenging and effortful, while avoiding rapid physical exhaustion as when asking participants to exert maximal force at each trial. Participants were not provided with any real-time feedback about the force they were applying during this phase to prevent them from using effort minimization strategies (i.e., keeping their force close to the threshold in order not to unnecessarily exceeding it too much). Once the force task ended, the go cue was replaced by a visual analogical scale on the screen and participants could stop applying any force on the dynamometer. The visual scale was represented as a gray horizontal line, ranging from 1 (left extremity) to 100 (right extremity), with a white bar superimposed vertically on the middle of the line. This white bar was a cursor that participants could slide along the horizontal line to rate how effortful they felt it was to apply enough force on the dynamometer during the preceding force task. They displaced the cursor using a computer mouse with their left hand (to avoid moving the right hand away from the dynamometer) until it was positioned over the desired score. Visual feedback about performance during the preceding force task was given only after the subjective rating of effort was completed to ensure that effort evaluation was not biased by it. This feedback consisted in a message which content varied according to the percentage of time spent above the force threshold during the force task (> 50%: “Well done!”, 25% to 50%: “Not bad!”, < 25\%: “Could do better!”).

#### 2. EXP2

Participants were asked to keep their right hand open until the go cue was displayed. They were instructed to close their hand 4 times as soon as the go cue appeared, with movements executed with maximal amplitude at each repetition. The go cue remained on screen for 8 s and was then replaced by a black screen indicating intertrial interval if it was a Comf trial, or by visual feedback about their performance at the motor task if it was a Fast or Slow trial. Feedback was positive (“Bravo!”) if the mean MT of the 4 repetitions of hand opening reached the speed criterion, and negative if the mean MT fell outside (“Too fast!” / “Too slow”). The aim of this feedback was to encourage participants to correct their MT if they started slowing down in Fast or speeding up in Slow. Speed criteria in Fast and Slow were determined based on the distribution of MT values in the first block of Comf trials. Fast speed criterion corresponded to mean MT – 3*standard deviation, implying that participants received positive feedback in Fast whenever their mean MT was equal or inferior to this criterion. In Slow, participants received positive feedback whenever their mean MT was equal or superior to mean MT + 3*standard deviation.

### Data processing

#### 1. Behavior

Data processing was conducted on Matlab R2023a (MathWorks). In EXP1, movement force was computed as mean force value during the motor task, in percentage of the MVC. MT was calculated as the delay between the first and second crossing of 3 kg of force values during the force task, and RT as the delay between the appearance of the go cue and the first time point when the force exerted on the dynamometer exceeded 3 kg. In EXP2, MT was computed as the time separating 2 peaks of luminosity detected by the light sensor, corresponding to the moments when the hand was fully open. MT was averaged across the 4 repetitions. RT was computed as the time separating go cue onset and the first luminosity peak. |Acc| corresponded to the mean absolute acceleration between go cue onset and the end of the fourth repetition. Outliers were calculated separately for each motor variable per participant and condition. They were defined as values inferior to the median minus 3 absolute deviations around the median (MADs)^50^ or superior to the median plus 3 MADs. They were removed from the analyses (4.6% and 2.6% trials in average across motor variables that have been measured in EXP1 and in EXP2, respectively).

#### 2. EEG

Preprocessing of raw EEG data was performed using EEGLAB toolbox in Matlab^56^. Similar EEG preprocessing pipelines were applied for data of EXP1 and EXP2. First, data was downsampled to 250 Hz. Then, a bandpass filter between 1 and 49 Hz was applied. Noisy channels were automatically detected using the Clean Raw Data algorithm from EEGLAB with the following parameters: signal flat for more than 5 s, standard deviation of high-frequency noise superior to 4, and correlation with nearby channels inferior to 0.8. The original signal from those channels was removed and interpolated based on the activity from their neighboring electrodes. Data was then segmented into epochs of 11.5 s duration locked around go cue (−11 s to +0.5 s). This period encompassed NF duration (2 to 10 s preceding go cue onset, see Neurofeedback section), with some additional time beforehand that included EEG activity at rest (i.e., during the preparatory period). Considering the minimum duration of one trial (∼16 s in EXP1 and 19 s in EXP2), the chosen time window length also prevented from overlapping between epochs. Independent component analysis (ICA) was run including epochs of all NF conditions for each participant separately using runica algorithm from EEGLAB. Artefactual components related to eye movements were identified based on their anterior location, spurious occurrences and low frequency dominant spectrum^57^. The signal was then re-referenced to the average scalp potential. Afterwards, time-frequency decomposition was performed with Morlet wavelets (5-40 Hz with 1 Hz step, 3 to 6.6 cycles (i.e., increment of 0.1 per 1 Hz)) using functions from FieldTrip toolbox^58^. Afterwards, mean β power was computed by averaging power values between 13 and 30 Hz during NF presentation at C3 electrode for EXP1 data, and D19 (C3 equivalent in ABCD system) electrode for EXP2 data, separately for each participant and experimental condition. β power was expressed as the percentage of change from baseline β power (i.e., β power = (actual β power – baseline β power) / baseline β power * 100). Considering that baseline β power that was used for NF calibration was acquired in different mental states in EXP1 (resting state) and EXP2 (3 s before go cue in the first block of Comf trials), baseline β power that was used for correcting β power in EXP1 was recomputed as mean β power during NF presentation of the first block of Sham-Passive trials. As for behavioral data, outliers were detected separately for each participant and condition and defined as values inferior to the median minus 3*MADs or superior to the median plus 3*MADs, before being removed from the analyses (3.9% and 2.4% of trials in average in EXP1 and EXP2, respectively). Outliers from the first blocks of each experiment were removed before computing baseline-corrected β power.

### Statistical analyses

Within-subject experimental designs were conducted in EXP1 and EXP2. Cluster-based permutation tests were first applied to β power from each experiment dataset separately, using functions from FieldTrip toolbox. NF conditions were compared to each other, in order to test if the signal from electrodes that were used to compute NF was significantly modulated across NF conditions, and if β power from other brain regions was impacted by NF. Then, a repeated measure ANOVA including data from EXP1 and EXP2 was conducted to compare β power across NF conditions (β-down, β-up and Sham-Passive). β power was also compared across speed instructions (Fast, Slow, Comf) in EXP2 data using repeated measure ANOVA. Greenhouse-Geisser’s correction was applied to p-values when sphericity assumption was violated (Mauchly’s test p-value < 0.05). Considering effective sample sizes (n = 28 in EXP1, n = 30 in EXP2), a medium effect size (f = 0.25) and a type 1 error set at 5%, the statistical power of this analysis was 81.2% in EXP1 and 84.1% in EXP2, based on calculations from G*Power 3^59^. Posthoc analysis with pairwise comparisons was conducted using paired Student’s t-tests when observations were normally distributed (Shapiro-Wilk’s test p-value ≥ 0.05), and Wilcoxon’s rank tests when normality assumption was violated (Shapiro-Wilk’s test p-value < 0.05). When comparing average value of a variable to a tested value, one-sample t-tests were used when observations were normally distributed (Shapiro-Wilk’s test p-value ≥ 0.05), and one-sample Wilcoxon’s rank tests when normality assumption was violated (Shapiro-Wilk’s test p-value < 0.05). P-values were corrected for multiple comparisons using the False Discovery Rate (FDR)^60^. Effect sizes were reported as partial eta squared (η²p) for ANOVAs, Cohen’s d (d) for paired Student’s t-tests, and rank biserial correlation (r) for Wilcoxon’s rank tests. Statistical analyses were conducted in Jamovi 2.3.28 and Matlab R2023a.

Changes in motor cortical β power and motor behavior were also assessed at the level of each participant individually, using paired t-tests on the distribution of single-trial data across NF conditions. Participants were classified as responders if they showed significant decreases in β power in β-down as compared to in β-up (i.e., p-value < 0.05 and t-value < 0), anti-responders if they demonstrated significant increases in β power in β-down as compared to in β-up (i.e., p-value < 0.05 and t-value > 0), and as non-responders if they demonstrated no significant change in β power when comparing β-down to β-up (i.e., p-value ≥ 0.05). Cohen’s d was reported as effect size.

LME models were conducted to determine the significance and direction of the relationship between each motor variable and β power, while considering variance due to individual differences. β power (continuous) and NF condition (3 levels: β-down, β-up, Sham-Passive) were included as fixed factors and Participant ID (1 level per participant) as random factor. The equation used for the model was the following: Motor variable ∼ 1 + β power + NF condition + β power: NF condition + (1 + β power + NF condition + β power:NF condition | Participant ID). An additional speed instruction factor (Fast, Slow) was included in LME models of motor variables of EXP2, as fixed factor and random slope, considering that the direction of the relationship between motor variables and β power was expected to potentially vary in function of speed instruction. Frequential specificity of the effect of β power on motor variables was assessed by conducting LME models including spectral power in a 1 Hz frequency band as β power factor (the equation used was the same as described above), separately for all frequencies between 8 and 35 Hz, motor variables and speed instruction in EXP2 (28 frequencies*3 motor variables = 84 models in EXP1; 28 frequencies*3 motor variables*2 speed instructions = 168 models in EXP2). Parameter estimates (slopes) of the effect of β power and their 95% confidence intervals were extracted from each model. In EXP1, only the first 10 trials of each block of active NF trials (β-down and β-up, n = 20 trials per block) were included in LME models to ensure adequate comparison of motor variables across NF conditions as mean force dropped significantly throughout each block (Figure S2, Table S1) and blocks of Sham-Passive trials comprised only 10 trials. Since no significant effect of block was found on MT in EXP2 (Figure S3, Table S2), all trials were included in LME models.

## Supporting information

Supplementary results

## Data availability

All data and computer code are available in a Zenodo repository through <u>this link</u> (doi: 10.5281/zenodo.14638353). Requests should be addressed to E.P.

## Acknowledgements

This work was supported by the French National Research Agency (BETAPARK Project, ANR-20-CE37-0012) and the University of Bordeaux IdEx “Investments for the Future” program / GPR BRAIN_2030.

## Author contributions

C.D., L.P. and E.P. designed the experiments (EXP1: C.D., L.P. and E.P.; EXP2: E.P.), E.P. and C.G. conducted the experiments (EXP1: E.P., EXP2: E.P. and C.G.), E.P. and A.P-V. analyzed the data (EXP1: E.P., EXP2: E.P. and A.P-V), E.P. wrote the paper and C.J.-K., N.G. and B.L. edited the paper.

## Materials and correspondence

Correspondence and materials requests should be addressed to E.P. at.

