## Supplementary results for "Changes in cortical beta power predict motor control flexibility, not vigor"

1. Changes in β power measured online across NF conditions and trials in EXP1 and EXP2

In EXP1 and EXP2, β power was estimated in (nearly) real time as the squared amplitude of the signal at C3 (EXP1) / D19 (EXP2) electrode between 13 and 30 Hz, averaged over 500-ms epochs with 50% overlap between them (see Methods). This procedure (simplified computation of signal power and smoothing) is commonly used to ensure accurate neurofeedback timing (no temporal lag because of time-consuming power computation) and readable neurofeedback content (no flickering because of variable β power values) in neurofeedback studies.^6,23,34^ This “online” measurement of β power represents a better proxy for NF performance than β power computed offline, as it faithfully reflects the level of the gauge.

In EXP1, online β power averaged across trials was significantly impacted by NF condition (F(2,54) = 12.7, p = 10^-5^, η²p = 0.32). Online β power was decreased in β-down as compared to Sham-Passive (t(28) = -3.8, p = 0.001, d = -0.71) and β-up (t(27) = -4.1, p = 0.001, d = -0.77), but no significant difference was found between Sham-Passive and β-up (t(27) = 1.5, p = 0.143, d = 0.29) (Figure S1A). At the trial level, online β power appeared decreased in β-down in comparison to β-up from the first block of trials (Figure S1B). Likewise, in EXP2, online β power averaged across trials was significantly impacted by NF condition (F(2,58) = 4.6, p = 0.014, η²p = 0.14). Online β power was reduced in β-down as compared to Sham-Passive (t(29) = -3.4, p = 0.006, d = -0.62) and there was a trend toward reduced β power in β-down in comparison to β-up (t(29) = -2.2, p = 0.052, d = -0.40), but no significant difference was found between Sham-Passive and β-up (t(29) = 1.1, p = 0.295, d = 0.20) (Figure S1C). Similarly to EXP1, at the trial level, online β power appeared decreased in β-down in comparison to β-up from the first block of trials (Figure S1D).

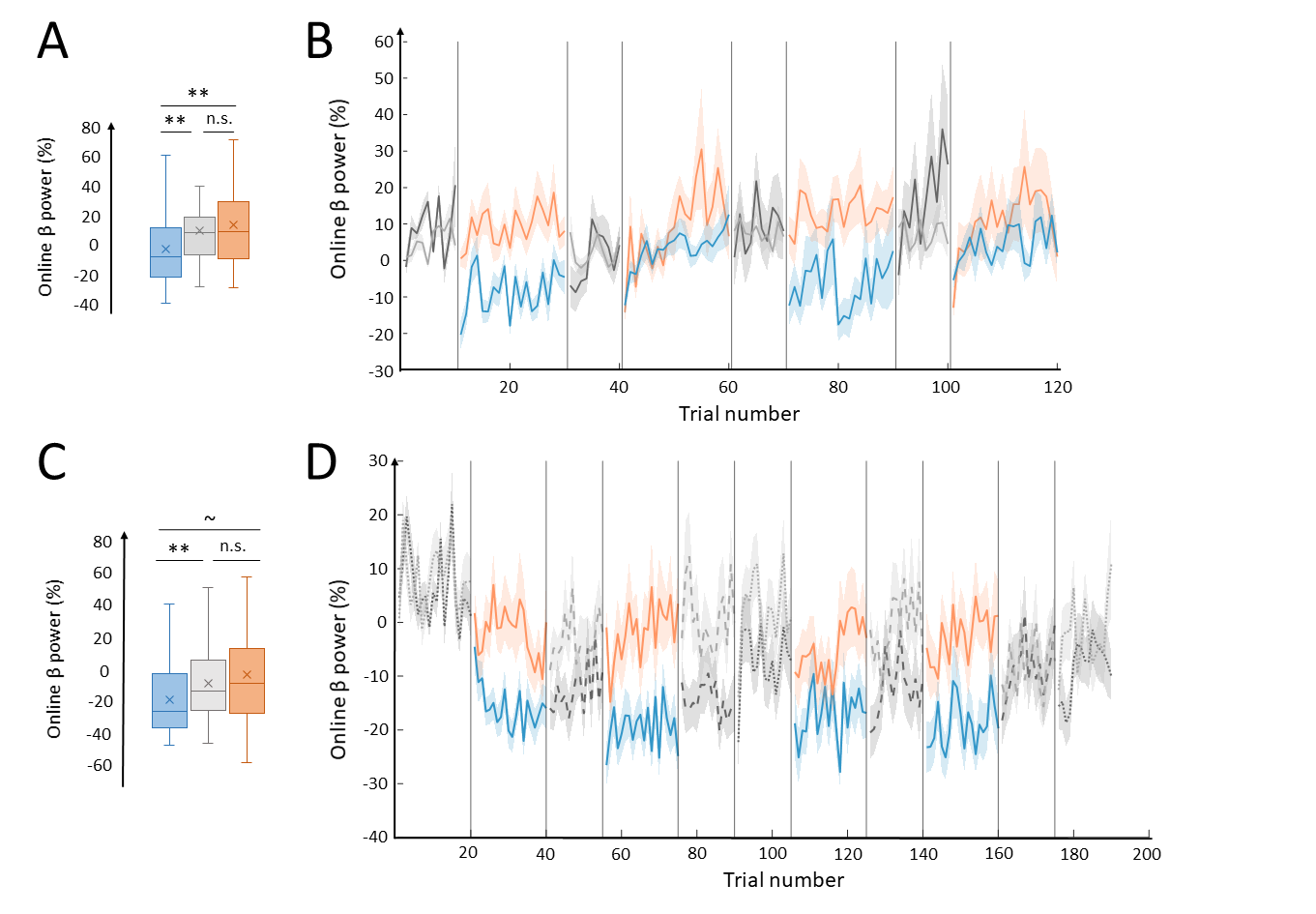

**Figure S1.** Online changes in β power across NF conditions. **A.** Average online β power in EXP1 in β-down (blue), Sham-passive (gray), and β-up (orange). Mean and median values of the distributions are respectively indicated by a cross and a horizontal line. **B.** Evolution of average online β power across trials in EXP1, represented in a chronological order. Blue and orange lines indicate β-down and β-up trials, respectively. Dark gray lines illustrate Sham-passive trials pertaining to participants who began with a block of β-down trials, and light gray lines represent Sham-passive trials pertaining to participants who began with a block of β-up trials. Vertical lines depict separation between blocks of trials. Shaded areas illustrate 95% confidence intervals. **C.** Average online β power in EXP2 in β-down (blue), Sham-passive (gray), and β-up (orange). Mean and median values of the distributions are respectively indicated by a cross and a horizontal line. **D.** Evolution of average online β power across trials in EXP2, represented in a chronological order. Blue and orange lines indicate β-down and β-up trials, respectively. Sham-passive trials are represented as dashed lines and Comfortable trials as dotted lines. Dark and light grey lines illustrate online β power in the β-down and β-up session, respectively. Vertical lines depict separation between blocks of trials. Shaded areas illustrate 95% confidence intervals. ***p < 0.001, **p < 0.01, *p < 0.05, ~ 0.05 < p < 0.1, n.s. = not significant.

2. Changes in force throughout trials in EXP1

As expected, once the go cue appeared on screen, mean force abruptly increased and was then maintained for a few seconds at a high level (Figure S2A). Mean force appeared to decrease across and within blocks (Figure S2B). Indeed, ANOVA including a Trial factor (1, 2, 3, 4 ; each corresponding to a series of 10 consecutive trials) and NF condition (β-down, β-up, Sham-Passive) showed a significant effect of Trial (F(1.5,41.7) = 23.7, p = 10^-6^). Posthoc analysis revealed significant differences in mean force across all Trial levels, except between 2 and 3, corresponding to the 10 last trials of the first block and 10 first trials of the second block in β-down and β-up (Table S1). No significant interaction was found between NF condition and Trial (F(4.1,110.0) = 1.2, p = 0.330), demonstrating that the significant effect of Trial on mean force did not significantly vary across NF conditions. This finding highlights a significant progressive decrease in mean force within and across blocks of trials.

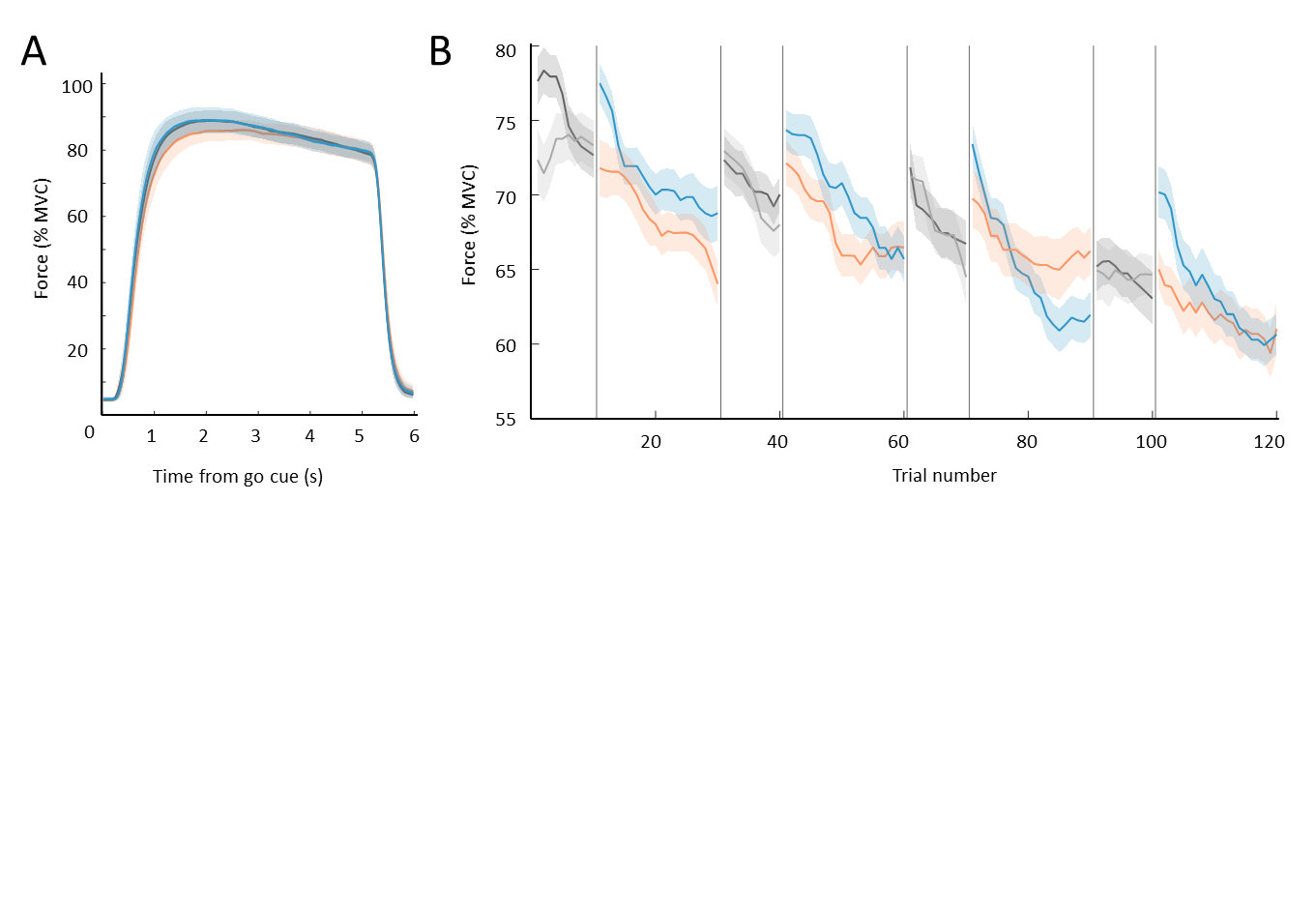

**Figure S2.** Changes in movement force within and across trials in EXP1. **A.** Movement force within trials, averaged over trials and participants. Mean movement force in β-down, β-up and Sham-Passive is highlighted in blue, orange and gray, respectively. Go cue appeared on screen at time 0 and disappeared 5 s later. **B.** Movement force across trials, averaged across participants who started with a block of β-down trials (n = 14) and those who started with a block of β-up trials (n = 14) separately. Mean force during blocks of Sham-Passive trials is illustrated in dark gray for participants who started with a block of β-down trials, and in light gray for participants who started with a block of β-up trials. Mean force in blocks of β-down and β-up trials is highlighted in blue and orange, respectively. Black vertical lines indicate separations between blocks.

| Statistics | T-value | FDR corrected p-value |
| --- | --- | --- |
| **Trial 1** (Block 1, 10 first trials) – **Trial 2** (Block 1, 10 last trials) | 5.6 | 10^-5^ |
| **Trial 1** (Block 1, 10 first trials) – **Trial 3** (Block 2, 10 first trials) | 4.1 | 10^-4^ |
| **Trial 1** (Block 1, 10 first trials) – **Trial 4** (Block 2, 10 last trials) | 6.4 | 10^-6^ |
| **Trial 2** (Block 1, 10 last trials) – **Trial 3** (Block 2, 10 first trials) | 1.2 | 0.230 |
| **Trial 2** (Block 1, 10 last trials) – **Trial 4** (Block 2, 10 last trials) | 4.3 | 10^-4^ |
| **Trial 3** (Block 2, 10 first trials) – **Trial 4** (Block 2, 10 last trials) | 6.7 | 10^-7^ |

**Table S1.** Results from posthoc analysis of Trial effect on mean force in EXP1.

3. Changes in MT throughout trials in EXP2

MT appeared strongly impacted by speed instruction, as demonstrated by modulations of the distance between two luminosity peaks recorded with the luxmeter placed on the palmar side of the right index finger (Figure S3A). MT appeared to modestly vary across and within blocks from a given experimental condition (Figure S3B). A 3-way ANOVA including a Trial factor (1, 2, 3, 4 ; each corresponding to a series of 10 consecutive trials), NF condition (β-down, β-up, Sham-Passive) and Speed (Fast, Slow) highlighted a significant main effect of Trial on MT (F(1.7,47.9) = 4.0, p = 0.031). However, posthoc analysis did not reveal any significant difference in MT across and within blocks after FDR correction (Table S2).

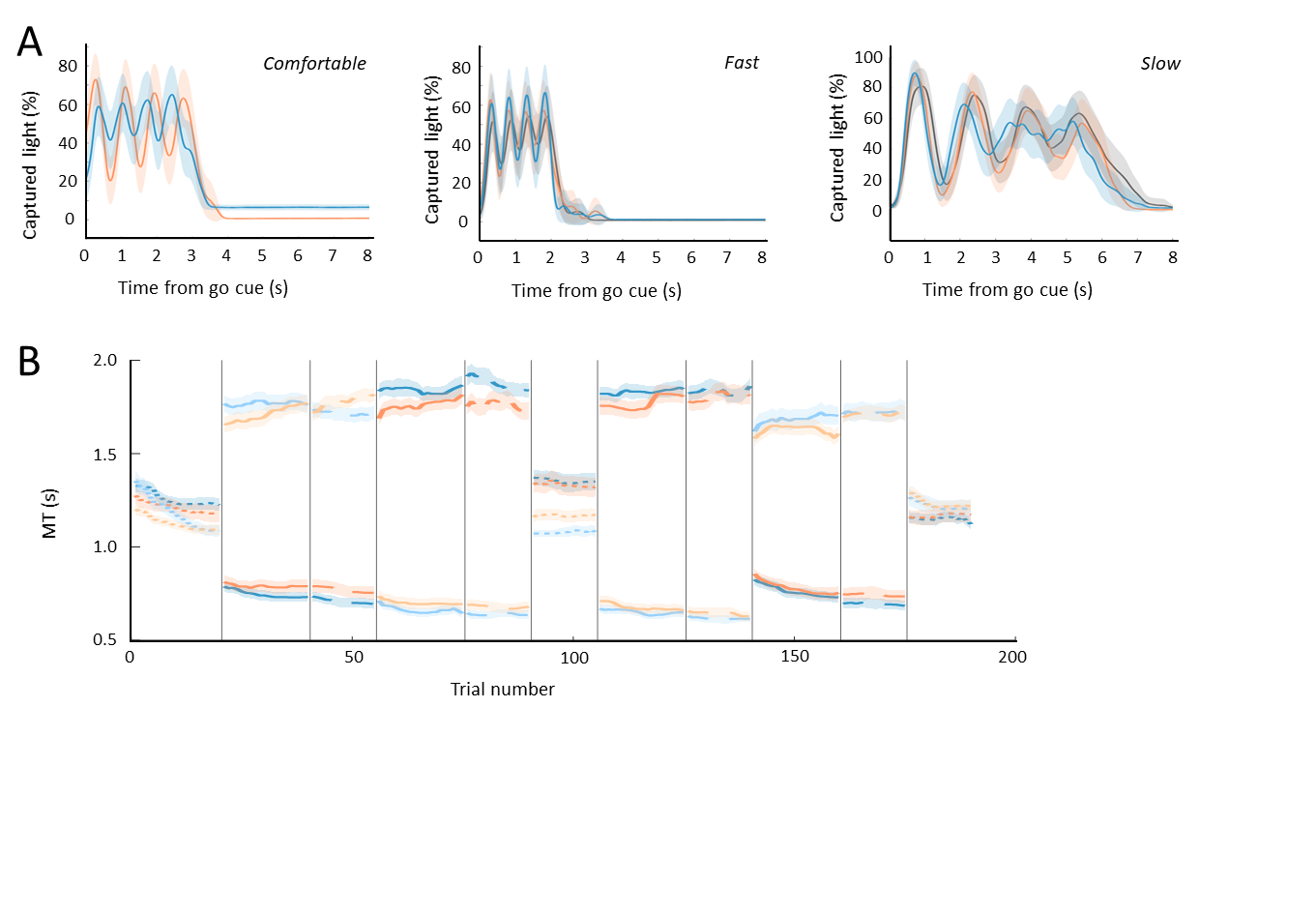

**Figure S3.** Changes in MT within and across trials in EXP2. **A.** Light captured by the luxmeter within trials, averaged over trials and participants in Comf (left), Fast (middle) and Slow (right). MT was computed as the average time between two luminosity peaks. Mean captured light in β-down, β-up and Sham-Passive is highlighted in blue, orange and gray, respectively. Go cue appeared on screen at time 0 and disappeared 5 s later. **B.** MT across trials, averaged across sessions and across participants who started with a session of β-down trials (n = 15) and those who started with a session of β-up trials (n = 15) separately. MT during β-down and β-down sessions is highlighted in blue and orange, respectively. Average MT of participants who started with the β-up session is represented in light blue and orange and, therefore, average MT of participants who started with the β-down session is represented in darker blue and orange. Dotted and dashed lines respectively indicate MT in blocks of Comf and Sham-Passive trials.

| Statistics | T-value | FDR corrected p-value |
| --- | --- | --- |
| **Trial 1** (Block 1, 10 first trials) – **Trial 2** (Block 1, 10 last trials) | 0.1 | 0.916 |
| **Trial 1** (Block 1, 10 first trials) – **Trial 3** (Block 2, 10 first trials) | 1.7 | 0.148 |
| **Trial 1** (Block 1, 10 first trials) – **Trial 4** (Block 2, 10 last trials) | 2.2 | 0.099 |
| **Trial 2** (Block 1, 10 last trials) – **Trial 3** (Block 2, 10 first trials) | 1.7 | 0.189 |
| **Trial 2** (Block 1, 10 last trials) – **Trial 4** (Block 2, 10 last trials) | 2.5 | 0.107 |
| **Trial 3** (Block 2, 10 first trials) – **Trial 4** (Block 2, 10 last trials) | 1.6 | 0.150 |

**Table S2.** Results from posthoc analysis of Trial effect on mean force in EXP2.
